# A novel classification framework for genome-wide association study of whole brain MRI images using deep learning

**DOI:** 10.1101/2024.01.11.575251

**Authors:** Shaojun Yu, Junjie Wu, Yumeng Shao, Deqiang Qiu, Zhaohui S. Qin, the Alzheimer’s Disease Neuroimaging Initiative

## Abstract

Genome-wide association studies (GWASs) have been widely applied in the neuroimaging field to discover genetic variants associated with brain-related traits. So far, almost all GWASs conducted in neuroimaging genetics are performed on univariate quantitative features summarized from brain images. On the other hand, powerful deep learning technologies have dramatically improved our ability to classify images. In this study, we proposed and implemented a novel machine learning strategy for systematically identifying genetic variants that lead to detectable nuances on Magnetic Resonance Images (MRI). For a specific single nucleotide polymorphism (SNP), if MRI images labeled by genotypes of this SNP can be reliably distinguished using machine learning, we then hypothesized that this SNP is likely to be associated with brain anatomy or function which is manifested in MRI brain images. We applied this strategy to a catalog of MRI image and genotype data collected by the Alzheimer’s Disease Neuroimaging Initiative (ADNI) consortium. From the results, we identified novel variants that show strong association to brain phenotypes.

## Introduction

Over the past decade, genome-wide association studies (GWASs) have been successfully applied to many diseases and traits and identified a large number of genotype-phenotype associations^1,2^. One notable success of GWAS is in the field of neuroimaging genetics, which aims to characterize, discover, and evaluate the association between genetic variations and brain imaging measurements^3^. Neuroimaging studies of twins and their siblings have revealed that numerous key brain structures are strongly influenced by genetic factors^4–11^. Recent advances in acquiring high-dimensional brain imaging and genome-wide data have provided unprecedented new opportunities to study the effect of genetic variations on a wide range of brain phenotypes. Exciting findings have been made using GWAS that connects genetic variants with brain functions and anatomical structures^3,12–20^.

Despite many novel findings, almost all neuroimaging GWASs conducted so far use pre-determined hand-crafted summary statistics, named imaging quantitative traits (iQTs) or imaging-derived phenotypes (IDPs)^1^, drawn from individual voxels^2^ or a pre-specified Regions-of-Interests (ROI) of the brain^3–5^ for univariate analyses. Such a practice might overlook important imaging characteristics that exist in full frame images, such as texture and subtle variations in the signal intensity of the images. Although methods have been developed to conduct GWAS on multiple traits simultaneously such as multifactor dimensionality reduction (MDR)-based and independent component analysis (ICA)^21^-based methods^6–9^, they are not designed to analyze high-dimensional structured phenotypes such as images. Furthermore, these methods only explore linear relationships, which is inadequate for analyzing complex phenotypes like Magnetic Resonance (MR) images. These limitations originate from the fact that GWAS as we know it is built under the hypothesis testing framework.

In this study, we present an innovative alternative strategy: conducting GWAS under a classification framework. For each single nucleotide polymorphism (SNP), we use the genotypes of this SNP as the label to divide all images into distinct subgroups, and then train a convolutional neural network (CNN)-based classifier to separate different subgroups of images. The hypothesis behind our strategy is that if an SNP is associated with brain-related traits, then there is high likelihood that detectable differences exist between brain images belonging to different genotype groups of this SNP. And such differences, albeit subtle, can be detected by adequately trained machine learning algorithms. Please see Figure 1.A for an illustration of our proposed strategy.

**Figure 1.**
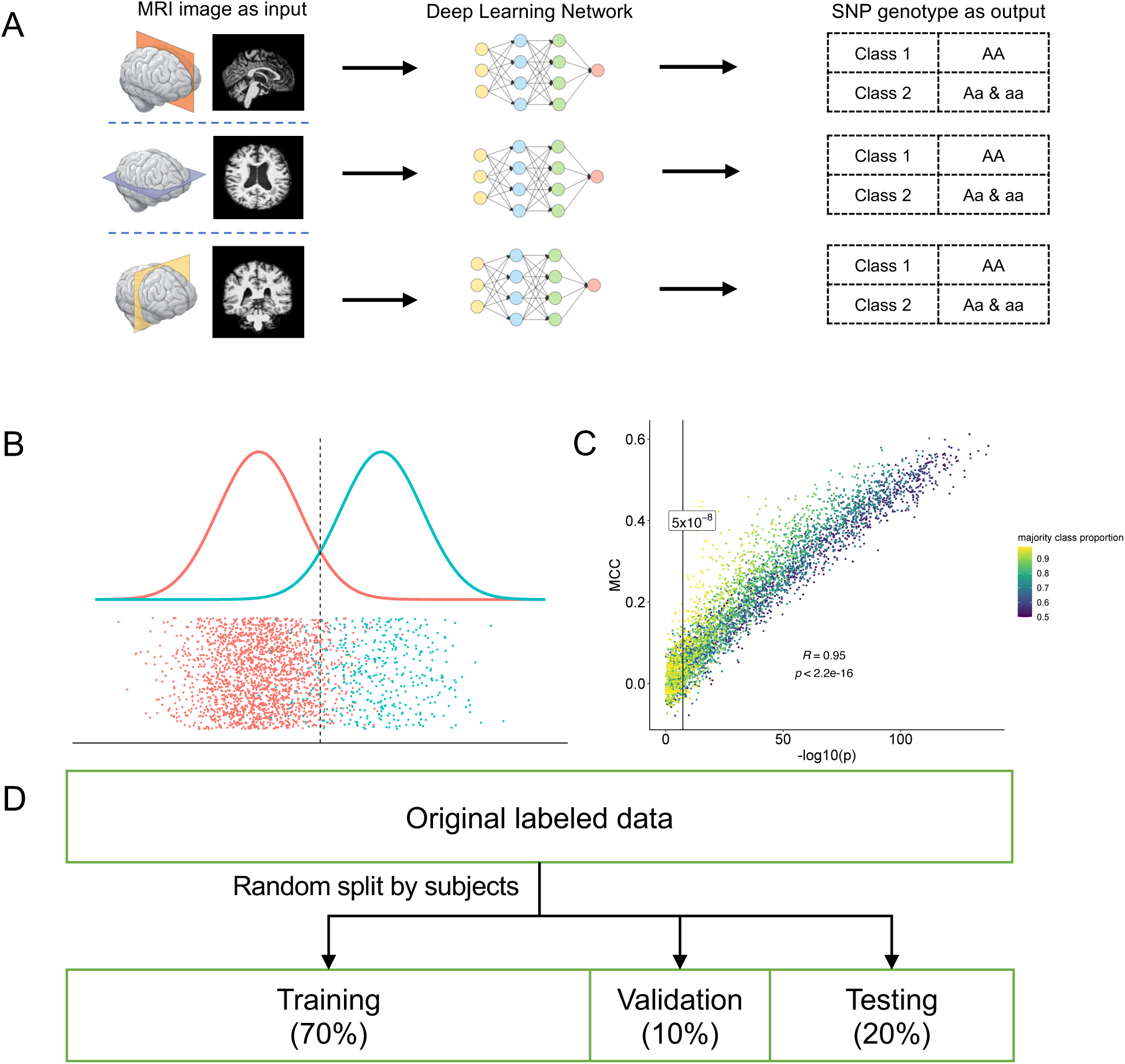
Illustration of the proposed classification-based GWAS framework. (A) Schematic representation of the proposed classification-based GWAS framework for conducting association study on Magnetic Resonance (MR) images, where full-frame brain MR images from different directions--axial, coronal, and sagittal, serve as input and SNP genotypes treated as labels. Classification was carried out by convolutional neural network (CNN) models. (B) Schematic illustration of the simulation study, where classification and testing are performed on the same dataset. Data are generated from two distinct univariate normal distributions with the same variance but different means. (C) Comparing p-values of two-sided independent t-test with classification performance metrics Matthew’s Correlation Coefficient (MCC) in the simulation study. (D) Illustration of evaluation process of the classification task in which all images are randomly divided into training, validation and testing sets by subjects in the ratio of 7:1:2.

A fundamental advantage of our new strategy is to enable association study to be conducted on high-dimensional phenotypes in a conceptually simple and natural way. Compared to the classical hypothesis testing-based strategy, our method relies on a different set of criteria to identify genotype-phenotype association. Under the hypothesis testing framework, the goal is to test whether a specific model parameter is the same across the two populations. P-values were used to represent the level of difference. In contrast, under the supervised classification framework, the focus is placed on finding ways to better distinguish samples belonging to the two populations. A completely different sets of performance measures were used.

Comparing to existing GWASs, our proposed method boasts two key novelties. First, our method can handle ultra-high-dimensional traits. As an example, in this study, the dimension of the three phenotypes we studied is either 33,124 or 39,676. Second, our method can accommodate non-linear relationships between genotypes and phenotypes. Both pose strenuous challenges under the traditional hypothesis testing framework. But under the classification framework, deep learning-based classifiers have been routinely utilized in computer vision and demonstrated extraordinary performance. We believe this powerful deep-learning-based method is able to make novel discoveries under a GWSA context.

## Results

### 1. Simulation Results

For the newly proposed strategy, it is of great interest and importance to understand how its result compares to the result obtained using the traditional hypothesis testing strategy, when applied to the same set of data. In order to understand the relationship between p-values and classification performance measures such as Matthew’s correlation coefficient (MCC) and macro F1 values, we conducted a series of simulation studies (Figure 1.B, Supplementary Figure 2, and Supplementary Table 1). For easy comparison, simulations were conducted on univariate variables. To be specific, random variables were sampled from two distinct Normal distributions with varying parameters. From the results, we found that overall, there is good agreement in terms of correlation between the p-values (from the hypothesis testing framework, −log() transformed) and the MCC or macro F1 values (from the classification framework) with a strong correlation ranging from 0.94 to 0.96 (spearman correlation coefficient, Figure 1.C and Supplementary Figure 2). We also notice that “moderate” classification measures ( 0.29≤ *MCC* ≤ 0.31) often correspond to highly significant testing results (p-values ranging from 10^-16^ to 10^-80^), especially when the sample size is large.

### 2. ADNI data

All the real data used in this study were obtained from the ADNI study (https://adni.loni.usc.edu/)^22^. The ADNI project recruited over 1,700 participants ranging from cognitively normal (CN) to patients with significant memory concerns (SMC), mild cognitive impairment (MCI), and Alzheimer’s disease (AD). ADNI aims to combine data from multiple imaging technologies, biomarkers, and clinical and neuropsychological assessment to study the progression of MCI and early AD. In this study, we primarily use the MRI and genotype data from ADNI 1 and ADNI 2. The detailed characteristics of the subjects are summarized in Supplementary Table 2. After quality control and preprocessing, we retain 13,722 2D MRI images and 313,267 SNPs from 1,009 subjects to be used in this study. Since ADNI is a longitudinal study, most participants contributed multiple MRI scans (Supplementary Figure 1.A). The demographic characteristics and the histogram of the data are presented in Supplementary Figure 1 and Supplementary Table 2, respectively.

### 3. Overview of the Method

We propose to conduct GWASs on full frame MR images of the brain under the classification framework. In classical GWAS, the phenotype, such as height, is treated as the response variable and genotypes are treated as explanatory variables. In our proposed method, genotypes are used to define classes (referred to as labels in the machine learning literature) and whole brain MR images are used as input. The underlying hypothesis posits that if a variant influences brain-related phenotypes, then different genotypes of the variant will exhibit distinct manifestations in the MR images. Such difference, albeit subtle, can be recognized by properly trained classifiers. In fact, different patterns in functional MRI have long been observed among patients with different genetic mutations such as the APOE variants^23^. Under the regression context, our new strategy effectively reverses the status of response and explanatory variables. In this study, to mitigate computation burden, we used 2D MR images, chosen to be the middle slice images from three different planes: axial, coronal and sagittal, derived from the original 3D T1-weighted (T1w) MR images. The details of the data processing can be found in the Methods Section. Given the 2D MR images, we trained CNN-based classifiers aiming to distinguish MR images with different genotype labels. Here we adopted a binary classification scheme, one genotype class consisted of samples with the homozygous wildtype genotype denoted as AA; and the other category contains samples with the heterozygous genotype denoted as Aa and homozygous mutant genotypes denoted aa. All CNN models used in this study shared the same architecture and complexity such that their performances are comparable. In the end, we rank the SNPs based on their classification performance.

For each SNP, after removing subjects with missing genotype information at the SNP, the MR image collection was split into three subsets: a training set, a validation set, and a test set with the ratio of 7:1:2 (Figure 1.D). The split is carried out in such a way that images from the same subject only appear in one of the three subsets to keep the three subsets independent. Then a CNN-based classification model was trained for a fixed number (30 in the present study) of epochs, and the best model during the training process was selected based on its performance measure on the validation set. During training, to save computing time, the process will be early stopped if there is no improvement in validation loss after ten epochs. Then, the final performance of the model was evaluated on the test set.

### 4. Model specification

Due to the fact that a massive number of classifiers need to be trained, we design our CNN model to be as simple as possible while maintaining descent performance. To help decide on the structure of the CNN model, we used the sex classification task as a benchmark. In addition to evaluating the models based on their classification performance, we also paid attention to the training time required to achieve this performance. We tested models with varying level of complexity and compared with state-of-the-art CNN models including ResNet^24^ and Xception^25^. Our results showed that modest CNN models produce excellent performance, with MCC 0.468, 0.387, and 0.619; macro F1 scores of 0.719, 0.682 and 0.805, in the axial, coronal, and sagittal planes, respectively, which is comparable to the state-of-the-art CNN models. Also, our modest CNN models required the least amount of time, at approximately 12 seconds to train for each task. In contrast, state-of-the-art CNN models such as ResNet and Xception require much longer training times, taking over 70 seconds to reach their best performance in training. These results are summarized in Supplementary Table 3. Overall, we found that modest CNN models can achieve comparable performance as the state-of-the-art CNN models while requiring significantly less training time, which makes them an attractive option for tasks where training time is a critical factor.

For each classification task, in order to account for uncertainty in the performance of individual CNN runs and minimize potential biases caused by confounding factors, we adopted a fine-tuning strategy. Details of the method is described in the Methods section. Essentially, we repeated CNN runs 20 times, and as comparison, we also performed permutation test 20 times and measured MCC and macro F1 scores each time. We then conducted two-sample t-tests to compare the two sets of performance measures and obtained p-values. Using sex as an example (Figure 2.A), we found that CNN models achieved significantly better performance in classifying sex-labeled MR images versus classifying label-permutated MR images with p-values 1.13×10^-48^, 4.76×10^-29^, 4.14×10^-53^, for the three planes respectively (Figure 2.B, C). These findings suggest that our CNN model is able to effectively capture features that distinguish MR images taken from males and females. In summary, our results demonstrate that the modest CNN model we adopted is able to successfully classify MR images with biologically meaningful labels such as sex.

**Figure 2.**
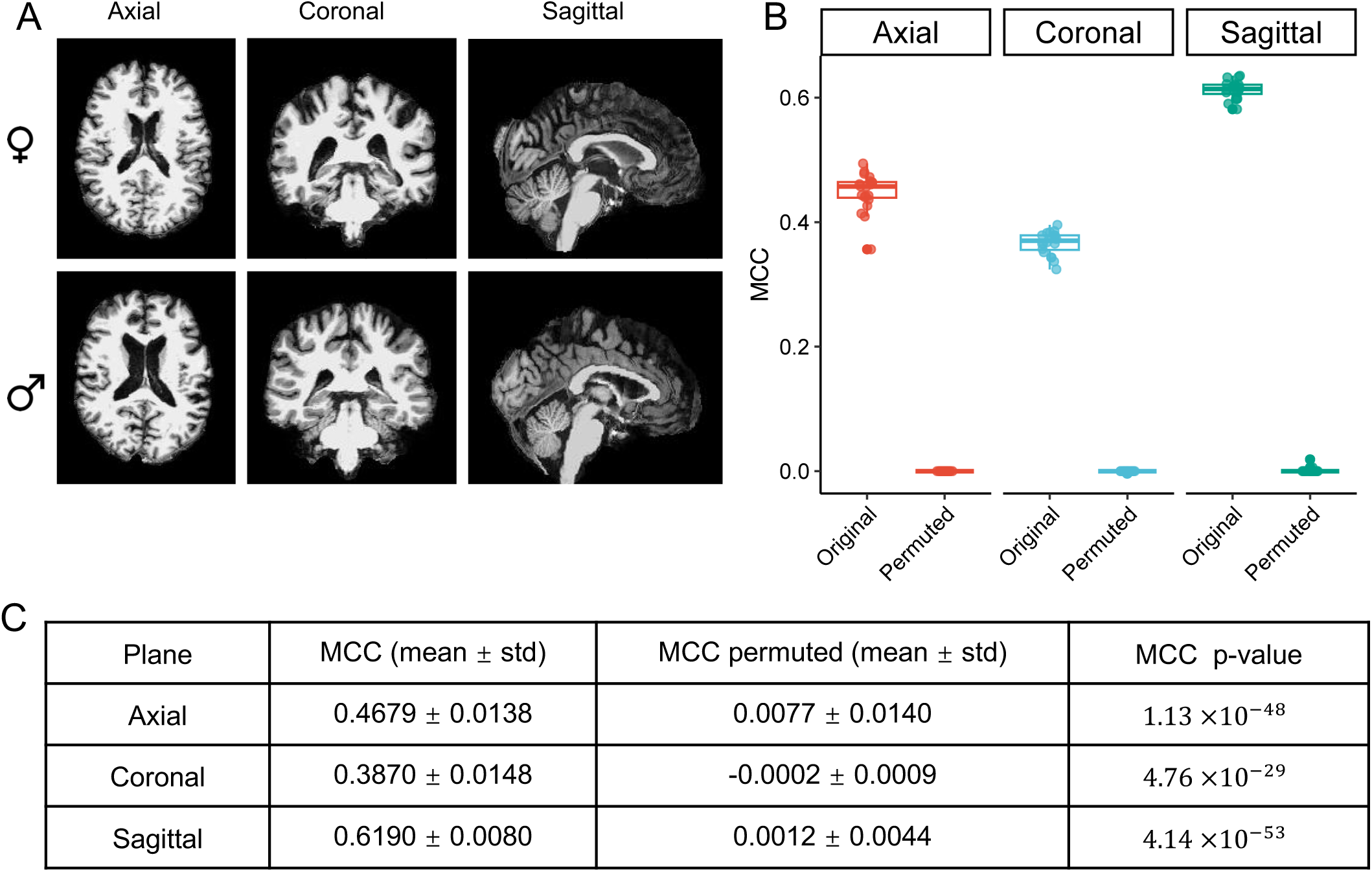
Results from sex classification using 2D full-frame MR images of the brain. (A) Sample 2D full-frame brain MR images from three different planes--axial, coronal, and sagittal. (B) Side-by-side boxplots showing MCC values from sex classification in axial, coronal, and sagittal planes before and after randomly permuting the sex labels. (C) Summary statistics and p-values from the fine tuning step of sex classification using 2D full-frame MR images of the brain.

### 5. GWAS results

Using MRI image slices from three different planes—axial, coronal and sagittal, we trained three different models for each SNP and measured their classification performance on the test set to identify potential associations between SNPs and brain MR images. Two different criteria were used: MCC and macro-F1 score. The performance results for all SNPs are presented in Supplementary Table 4. A histogram was constructed to visualize the distribution of MCC scores for all SNPs in different planes (Figure 3.A). The histogram showed that the distribution of MCC values was approximately normal with mean around 0, meaning that for most SNPs, genotype-labeled full-frame brain MRI images cannot be well classified using CNN-based classifiers, which is expected.

**Figure 3.**
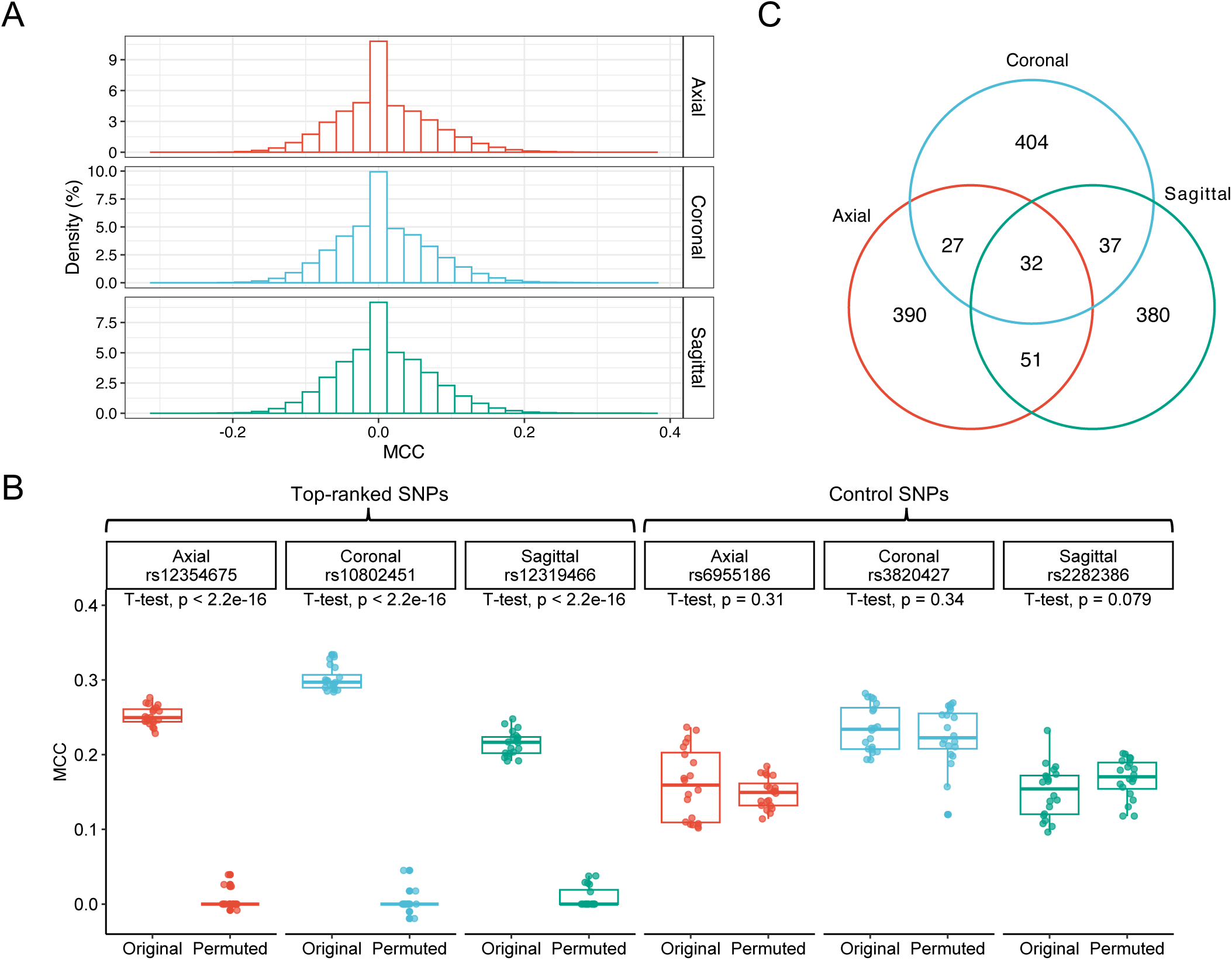
Main results from classification-based GWAS applied to Alzheimer’s Disease Neuroimaging Initiative (ADNI) MR image data. (A) Histograms of all MCC values derived from the GWASs conducted in each of the axial, coronal, and sagittal planes. (B) Illustration of the difference in the side-by-side boxplots between top-ranked SNPs and randomly selected low-ranked SNPs obtained from the GWAS conducted on three different planes--axial, coronal, and sagittal. The boxplot shows MCC values for classifying genotypes of the SNP, before and after randomly permuting the genotype labels. (C) Venn diagram displaying the number of overlapping SNPs among the three lists of top 500 SNPs ranked by MCC in the in the axial, coronal, and sagittal planes.

Using the MCC criterion, we selected the top 500 SNPs with the highest MCC values in each of the three different planes, which corresponds to top 0.018 percentile. From the Venn’s diagram (Figure 3.C) We saw that most of the SNPs are unique (with only 32 SNPs appear in all three lists), meaning using 2D images derived from different planes lead to unique association findings. Among the three lists of top 500 SNPs, 1,321 are unique (referred to as the 1,321 top SNPs hereafter). To select the very top among them, we apply the fine-tuning strategy, and rank these SNPs according to the two-sample t-test p-values. Figure 3.B shows examples comparing top-ranked and control SNPs in three different planes. Table 1 summarizes the top 20 SNPs ranked by the fine-tuning p-values.

**Table 1.** 20 Top SNPs significantly associated with full-frame MR image of the brain.

| SNP | Plane | MCC | MCC<br>P-value | Context | Gene | Phenotypes |
| --- | --- | --- | --- | --- | --- | --- |
| rs12354675 | Axial | 0.25 | $2.38 \times 10^{-40}$ | upstream gene<br>variant | *NPFFR1* | Depression;<br>**Right superior parietal** |
| rs10802451 | Coronal | 0.30 | $4.06 \times 10^{-39}$ | intron variant | *AHCTF1* | Depression;<br>**Mean Pallidum** |
| rs998461 | Axial | 0.22 | $3.26 \times 10^{-37}$ | regulatory region<br>variant | \ | Insomnia |
| rs2253927 | Coronal | 0.24 | $7.88 \times 10^{-37}$ | intergenic variant | \ | Alcohol dependency;<br>**Cerebellar vermal lobules** |
| rs1198414 | Axial | 0.35 | $2.30 \times 10^{-36}$ | intergenic variant | \ | Body Mass Index |
| rs6838213 | Axial | 0.33 | $9.73 \times 10^{-35}$ | intergenic variant | \ | Neuroticism; |
| rs3812663 | Axial | 0.26 | $2.43 \times 10^{-34}$ | intergenic variant | \ | Epilepsy;<br>Alcohol dependency |
| rs17102716 | Coronal | 0.23 | $1.51 \times 10^{-33}$ | intron variant | *CTC-367F4.1* | **Right pericalcarine;**<br>Insomnia |
| rs7768422 | Axial | 0.33 | $1.56 \times 10^{-33}$ | intron variant | *SLC22A16* | **Left isthmus cingulate;**<br>Autism spectrum disorder |
| rs12319466 | Sagittal | 0.22 | $1.91 \times 10^{-33}$ | non-coding<br>transcript variant | *RP11-588H23.3* | Cognitive;<br>Schizophrenia |
| rs981684 | Coronal | 0.21 | $2.11 \times 10^{-33}$ | missense variant | *TMEM99* | **Right caudal middle**<br>**frontal** |
| rs16903930 | Axial | 0.26 | $3.01 \times 10^{-33}$ | intron variant | *EGFLAM* | **Left pars triangularis** |
| rs2626192 | Axial | 0.30 | $3.29 \times 10^{-33}$ | regulatory region<br>variant | *LINC01193* | \ |
| rs6703902 | Coronal | 0.22 | $7.10 \times 10^{-33}$ | intergenic variant | \ | Neuroticism;<br>**Right rostral middle**<br>**frontal** |
| rs4895172 | Sagittal | 0.24 | $7.21 \times 10^{-33}$ | intron variant | *CTC-546K23.1* | Worrier / anxious feelings;<br>**Right accumbens area** |
| rs596985 | Coronal | 0.38 | $1.08 \times 10^{-32}$ | intron variant | *ROR1* | Depression; Cognitive |
| rs11845184 | Axial | 0.34 | $1.84 \times 10^{-32}$ | intron variant | *TSHR* | Neuroticism |
| rs11746806 | Coronal | 0.32 | $2.43 \times 10^{-32}$ | non-coding<br>transcript variant | *RP11-1023L17.1* | Hair color |
| rs7982701 | Coronal | 0.25 | $3.60 \times 10^{-32}$ | intron variant | *ZMYM2* | Cognitive;<br>**Mean Putamen** |
| rs4242182 | Axial | 0.29 | $1.12 \times 10^{-21}$ | intron variant | *MSX2* | Hair color;<br>**Right superior frontal** |

When using the macro F1 score, 1,301 of the top 500 SNPs identified across the three planes were unique. 957 SNPs appeared in the top 500 SNP lists ranked by both MCC and F1 (Supplementary Table 5, 6, 7). Supplementary Figure 3 presents the top SNPs ranked by macro F1 and their annotations.

#### Known associations of the top SNPs

To understand the properties of the top SNPs identified, it is of great interest to learn what traits they are associated with according to existing GWASs. To find out, we queried the top SNPs using the GWASATLAS^26^ tool and the detailed annotation results are presented in Supplementary Table 8, 9 and 10. Among the 1,321 top SNPs, we found that 859 (65.0 %) of them were associated with one or more known phenotypes at P < 0.01. Among the known phenotypes that these SNPs are linked to, many are neurological, psychiatric, or developmental-related such as depression, neuroticism, epilepsy, autism spectrum disorder and schizophrenia (Figure 4.A). It is particularly noteworthy that 11 out of the top 20 SNPs (55%) were already identified in previous studies as being associated with certain brain iQTs (bold text in Table 1) such as right superior parietal and mean pallidum.

**Figure 4.**
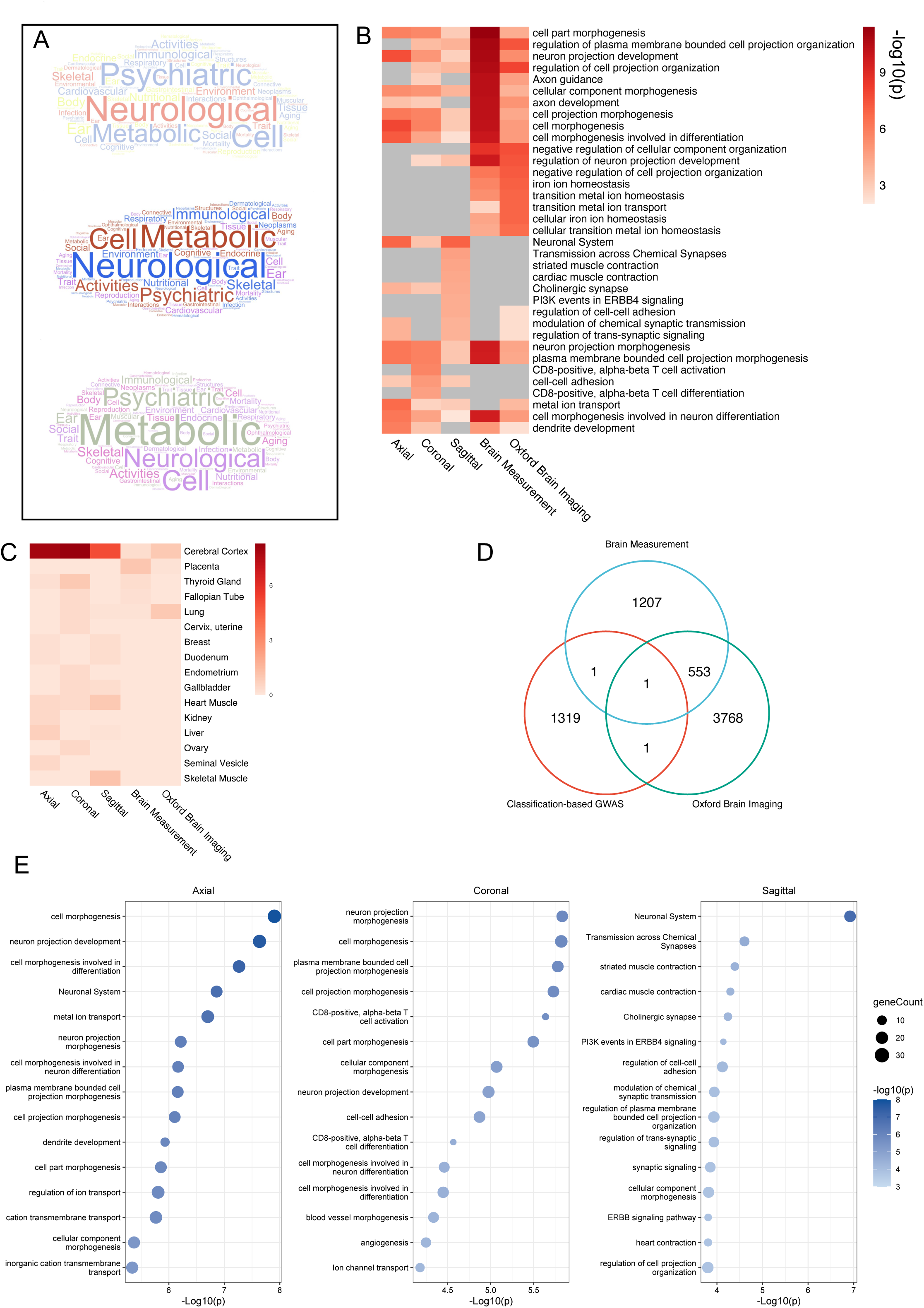
Exploring biological properties of genes close to the top SNPs. (A) A word cloud showing the major domains of phenotypes show up in PheWAS analyses of the 500 top-ranked SNPs identified from three different planes (from top to bottom): axial, coronal and sagittal. (B) The heatmap showing the enrichment levels (measured by −log transformed p-values) of GO terms and pathways from five lists of top-ranked genes. The first three lists correspond to top genes from our studies of classifying full-frame brain MR images from three different planes--axial, coronal, and sagittal. The fourth set was sourced from the GWAS catalog, the fifth one was acquired from the Oxford Brain Imaging Genetics (BIG) Server. Top 10 most enriched pathways identified from each of the five gene lists were used to generate this heatmap. A grey color indicates that the p-value is not significant (p > 0.05). (C) A heatmap comparing the enrichment (measured by −log transformed p-values) of the expression levels of the same five lists of top-ranked genes from a panel of 16 tissues using the TissueEnrich tool. (D) Venn diagram showing overlaps among top SNPs from three different sources: proposed classification-based GWAS, Brain Measurement (GWAS Catalog, 1762 SNPs) and the Oxford BIG (4323 SNPs). (E) Gene Ontology (GO) term and pathway enrichment analyses results from the GWASs conducted in each of the axial, coronal, and sagittal planes.

Further exploration into the genetic functions revealed that genes proximal to most of these 20 SNPs, have been previously highlighted in research for their substantial roles in brain and neurons. For example, the SNP with the most significant p-value is upstream of *NPFFR1*, also known as *GPR147*, which is a G protein-coupled Neuropeptide FF Receptor belonging to the RFamide receptor family. NPFFR1 plays a pivotal role in modulating stress and anxiety responses. In a recent study, Shen et al. found that the expression of the *Npffr1* gene is up-regulated in mice exposed to chronic unpredictable mild stress (CUMS) that exhibited depression-related behaviors^27^.

In another example, RP11-588H23.3, is a long non-coding RNA where the 10^th^ most significant SNP is located within. In a recent study, Gudenas and Wang found that the expression of RP11-588H23.3 is highly correlated with *CA2*, a key gene associated with intellectual disability^28^.

Another SNP on the list, rs11845184, is located in the intron of the *TSHR* gene, which encodes the receptor for the thyroid-stimulating hormone (TSH). TSHR plays an important role in the formation of neuroticism and aggression development^29^. It is also implicated in the actions of thyroid hormones, which are indispensable for normal brain development. These hormones facilitate neurogenesis, synaptogenesis, and myelination, as well as the differentiation and migration of neuronal and glial cells^30^. GWAS Catalog^31^ showed that this gene is associated with Tourette syndrome^32^ and Schizophrenia^33^.

Other SNPs also show interesting characteristics. For example, SNP rs16903930 is located in the intron of a gene named *EGFLAM* (EGF like, fibronectin type III and laminin G domains). GWAS Catalog showed that this gene is associated with multiple traits of interest including height^34^, mathematical ability^35^ and education attainment^36^. SNP rs596985 is located in the intron of a gene named *ROR1* (receptor tyrosine kinase like orphan receptor 1). GWAS Catalog showed that this gene is associated with multiple brain measurements^37^, cortical surface area^38^ and cortical thickness^38^. SNP rs7982701 is located upstream of a gene named *ZMYM2* (zinc finger MYM-type containing 2) which is shown to be associated with morphology-related traits such as body mass index^39^ and adult body size^40^ according to GWAS Catalog. SNP rs4242182 is located in the intron of *MSX2* (msh homeobox 2). GWAS Catalog showed that this gene is associated with cortical surface area^38^, posterior cortical atrophy and Alzheimer’s disease^41^, vertex-wise sulcal depth and brain morphology^38^.

#### Pathway enrichment analysis

To understand which pathways the 1,321 top SNPs significantly affect, we located genes overlay or next to these SNPs and then conducted gene-based enrichment analyses using pathway enrichment tool Metascape^42^, we found that several brain- and development-related functional categories are significantly overrepresented among these genes. These categories include the neuron system, neurogenesis, and cellular processes such as neuronal cell proliferation, differentiation, and maturation as well as cell adhesion (Figure 4.B & E). These pathways are known to play critical roles in cellular growth, and differentiation in brain development.

#### Tissue enrichment analysis

To explore in which tissue the top genes are highly expressed, we conducted tissue enrichment analysis using TissueEnrich^43^. Remarkably, our results revealed that collectively, the 1,321 top genes express the highest in cerebral cortex, the tissue most related to brain in the 16-tissue panel (Figure 4.C). Analysis using Enrichr^44^ showed that brain tissues from individual donors are the most enriched among all tissue in the GTEx collection (Supplementary Figure 4). The enrichment results again confirm remarkably intimate relationships between top SNPs and brain functions and activities.

#### Compared to traditional GWAS on iQTs

For our new method, it is of interest to compare our results with those obtained using traditional neuroimaging GWASs, so that we can put our findings in context. To do this, we performed enrichment analyses on two sets of SNPs that were obtained from published neuroimaging GWAS conducted on iQTs. The first set was sourced from the GWAS catalog, the second was acquired from the Oxford Brain Imaging Genetics Server^20^. From each data source, we adopted a similar strategy by selecting the top 500 SNPs with significant associations to brain MR imaging measurements and mapping them to the nearest genes.

Among the five sets of SNPs obtained using various methods, we found that while the GWAS catalog and Oxford brain imaging SNP lists overlap substantially, there is little overlap between each of the three SNP lists (from three different planes) obtained by our method and the two aforementioned SNP sets discovered by conventional neuroimaging GWASs. This indicates that our method is able to discover novel brain-SNP associations that may be missed by conventional GWAS (Figure 4.D). For pathway enrichment analyses, our results showed that all five sets of top genes showed significant and comparable enrichment in brain and neuron-related pathways. However, the level of enrichment in terms of tissue-specific expression showed that the three sets of top genes identified by our method is much higher in the cerebral cortex, the tissue most related to brain in the tissue panel, than the other two sets (Figure 4.B). These findings provide further support for the conclusion that the SNPs identified by our novel GWAS strategy are highly relevant to the brain and likely to be involved in brain-related traits or diseases.

### 6. Saliency maps

In computer vision literature, saliency map has been recognized as a powerful tool of visualizing attention, showing which parts of the input image contribute the most to the model’s prediction in CNN models^45^. Here we plan to use saliency map to help us understand which parts of the brain the SNP is associated with. This is because highlighted regions in the saliency maps indicate brain regions that are highly relevant to the classification task. We used GradCAM++^46^ to visualize the attention over input images. A visual explanation for the corresponding class label is generated by combining the positive partial derivatives of feature maps of the last convolutional layer with respect to a specific class label. Model weights saved during training were loaded again to generate saliency maps for each of the two classes. Here we present the saliency maps for sex and the genotype of the SNP rs11845184 (Figure 5). Highlighted areas in the first two saliency maps indicate the brain regions that closely related to the class label (sex or SNP genotypes). The third column shows the absolute difference between the saliency maps of the two labels, thereby allowing us to identify the areas that are most discriminative between the two classes. Such saliency maps can be applied to other SNPs to visualize the brain regions that are most relevant to genotypes of the SNP.

**Figure 5.**
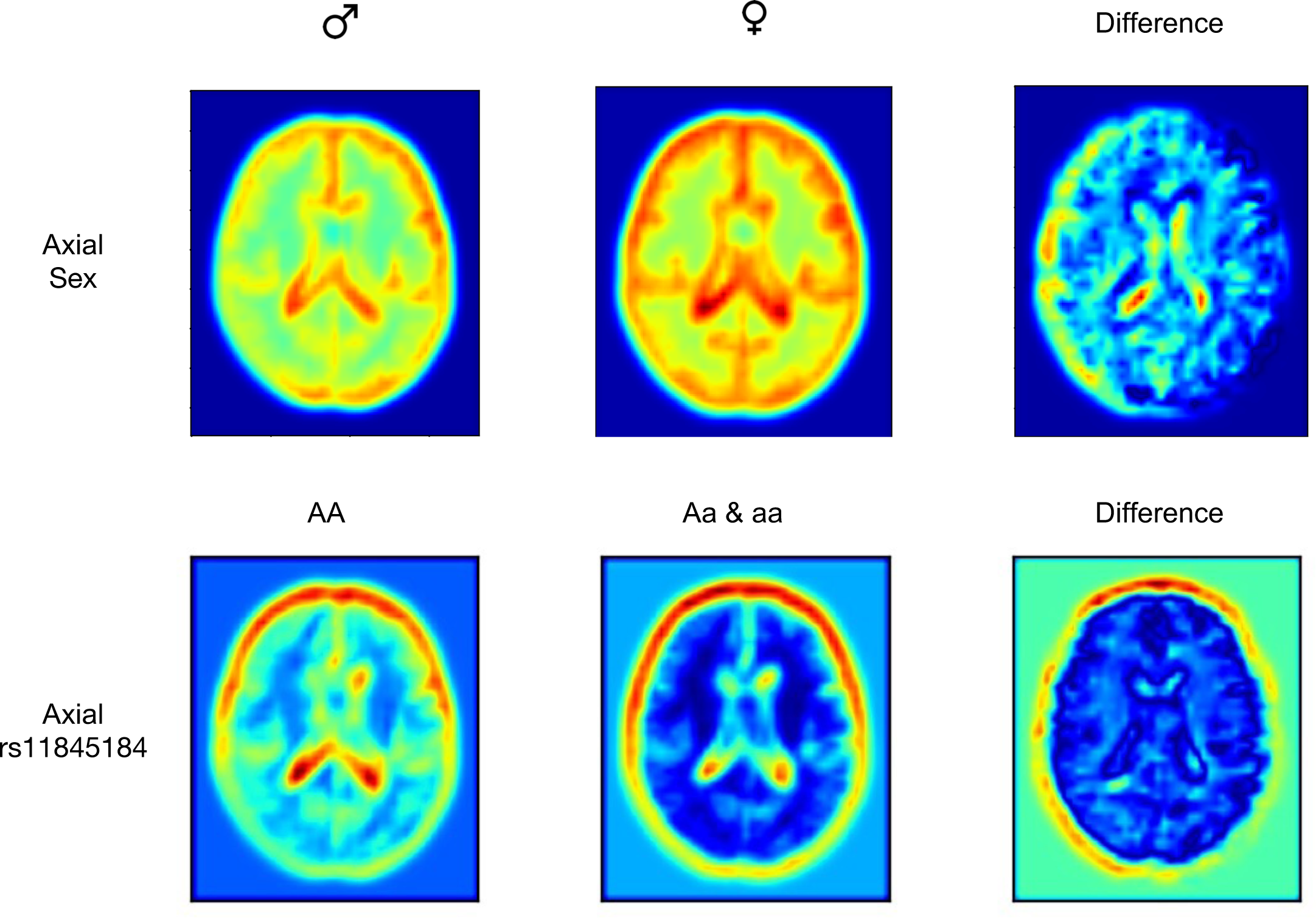
Saliency maps for the sex classification task and for the rs11845184 genotype classification task in the axial plane. The leftmost column shows the saliency map for male individuals, the middle column for female individuals, and the rightmost column for the difference between the two. The saliency maps in the first column depict the regions that are most important for the model to identify a male individual, while the second column illustrates the regions that are most important for the model to identify a female individual. The third column highlights the regions that are the most different between the two sexes, providing insight into the specific brain regions that are most differentiable.

## Methods

### 1. Performance evaluation measures

Our novel strategy uses well established classification performance measures to represent the association between full-frame MR images and SNP genotypes. Two metrics were selected to evaluate the performance: MCC and macro F1 score. These two metrics are known to provide robust measure of the overall classification performance, even with imbalanced datasets. This is important because for many SNPs, genotypes contain the alternative allele has low proportion^47^. The range of MCC is between −1 and +1. A coefficient of +1 represents a perfect prediction, random guesses produce MCC around 0. Let *tp, tn, fp* and *fn* represent the number of true positives, true negatives, false positives and false negatives, respectively. the MCC is defined as:

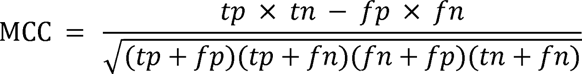

The F1 score is the harmonic mean of the precision and recall, defined as:

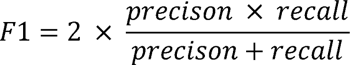

The range of the F1 score is between 0 and 1, where 1 indicates perfect classification and 0 indicates either precision or recall is 0. The relative contribution of precision and recall to the F1 score is the same.

For our classification tasks, there is no clear distinction between the positive label and the negative label, and such labels are required to calculate the precision and recall. Therefore, we use the macro F1 score instead. The macro F1 score calculates an F1 score for either label designation and then use their unweighted mean as the final result.

### 2. Simulation Study

In this study, we propose a novel classification-based strategy to conduct GWAS on high-dimensional phenotypes. Compared to traditional GWAS, we use classification metrics such as MCC and macro F1 score to measure the strength of association, which is different from the metrics used in traditional GWAS, such as association test p-values, therefore it is important to understand how the two sets of performance measures compared to each other. To achieve this, we conducted simulation studies in which we applied both classification and hypothesis testing on the same dataset and compared their performance measures. To be specific, for each simulated dataset, univariate data points were randomly drawn from two Normal distributions: *N* (0, 1) and N (μ, 1), where μ is fixed with values satisfy 0.2 ≤ μ ≤ 2, representing two different classes. For hypothesis testing, we used p-values from two-sample t-test to assess statistical significance. For classification, we used decision tree from R package rpart and compute the MCC and macro F1 score as performance measures. The simulation process is performed under different major class proportions (ranging from 0.5 to 0.98) and different values of μ. Since the size of *N* would affect the p-value, we also conducted this simulation study under different values of *N*, ranging from 3000 to 7000.

### 3. Genotyping data and processing

Genotyping for 1,009 participants were performed by investigators of the ADNI consortium using the Illumina Human 610-Quad, HumanOmni Express and HumanOmni 2.5 M BeadChips. PLINK was used to perform the standard quality control procedures for GWAS data. Excluding criteria include SNP call rate < 95%, Hardy-Weinberg equilibrium test p-value < 1 × 10^−6^, minor allele frequency (MAF) < 5%. Additionally, only SNPs located on the autosomes are included in the present study.

### 4. Neuroimaging data and processing

Original MR image data used in the study are 3D T1-weighted images from ADNI 1 and 2. To prevent the model from inadvertently learning from non-brain regions, all MR images were skull-stripped using the bet tool from the FSL software package^48^ (Oxford University, Oxford, UK). To account for variabilities in subject positioning and prescription of field-of-view, all high-resolution 3D T1-weighted images are normalized to the Montreal Neurological Institute (MNI) space^49^. To reduce the requirement for GPU memory and computational power, the images are resampled to 218 × 182 × 182 dimensions. To save computing time, we extract the middle slice from three different anatomical directions: sagittal, coronal, and axial using med2image (v.2.1.10, https://github.com/FNNDSC/med2image). The extracted images are in grayscale and standardized to be within 0 and 1.

### 5. CNN Model architecture and model training

Each of the preprocessed slices from the three directions was fed into a neural network separately. We used LeNet^50^, a simple CNN model structure, as the backbone with a few modifications: The model starts with a Conv2D layer with 8 filters, a kernel size of 3×3, and ReLU activation function. This is followed by a MaxPooling2D layer with a pool size of 2×2. The same pattern is repeated twice more with Conv2D layers with 16 and 32 filters, respectively, and additional MaxPooling2D layers. The output of the final MaxPooling2D layer is then flattened into a 1D array using a Flatten layer. Next, a Dense layer with 128 units and ReLU activation function is applied, followed by a Dropout layer with a rate of 0.5 to reduce overfitting. Finally, a Dense layer with 2 units and softmax activation function is used for classification. LeNet is one of the first CNN models. Although very simple, LeNet has been shown to achieve very good performance with low computational cost. Since we need to train over 900,000 models in total, it’s very important to design the model as simple as we possibly can while maintaining descent performance. For each SNP, we trained three induvial models using images from three different directions. The dataset for each binary classification task was split into three subsets: training, validation and test set with the ratio of 7:1:2. All the models were trained on a server with two TITAN RTX GPU cards. The average training time for an CNN model is about 20 seconds. Each model was trained at the max epochs of 30 and at a learning rate of 1e-4. The best model during training was selected based on the performance on the validation set and the weights of the best model were saved and used to evaluate the performance on the test set.

All models were developed based on the TensorFlow framework v2.4.3 and pre-trained weight on ImageNet is provided by Keras framework v2.4.0.

### 6. Permutation test of the classification performance

Performance measures such as MCC and macro F1 score are designed for a single classification task. But under the GWAS setting, the goal is to find out among hundreds of thousands of tasks, which ones are *classifiable*, meaning that there is substantial and recognizable difference exist between the two genotype-labeled classes. Given that deep learning models are complex and dynamic, its performance may vary when the model is retrained. To mitigate the uncertainty, we developed a fine-tuning strategy.

The procedure has three steps. First, for each SNP, using the original training, validation and test set, we retrain the CNN model 20 times, providing a distribution of classification performance results. Next, we randomly shuffle the labels of the images, and train a CNN model with the same architecture on the shuffled data. This process is repeated 20 times, providing a null distribution of the classification performance results. The third step is to test whether the classification performance on the original data is better than that of the shuffled data by comparing the two sets of performance measures using a one-sided two-sample t-test. This procedure is performed for both MCC and macro F1 score. Due to heavy computing cost, we only applied the fine-tuning strategy to a small fraction of top-ranked SNPs.

### 7. Pathway, tissue enrichment for genes

We biological pathways enrichment analysis on Gene Ontology^51^ biological process, Reactome Gene Sets^52^ and KEGG Pathway^53^ using Metascape^42^(version 3.5, http://metascape.org), and the results were visualized with the ggplot2 R package. Gene expression tissue enrichment was performed using TissueEnrich^43^ with human protein atlas data and Enrichr^44^ with GTEx project data (https://gtexportal.org/home/). TissueEnrich and Enrichr are web-based tools that utilize a hypergeometric test to calculate the enrichment of a series of curated and pre-defined gene sets within a list of input genes.

### 8. Querying related databases

We used a series of databases to annotate the top SNPs and their putative target genes. For the context and gene information, we used the Ensembl Variant Effect Predictor^54^. The Gene expression data were taken from the GTEx Project. In order to identify previous association studies for each independent locus, the genomic region contains these SNPs was first mapped to the hg38 version of the human genome using LiftOver, followed by a query against the GWAS Catalog database (https://www.ebi.ac.uk/gwas/) and GWAS atlas (https://atlas.ctglab.nl/)^26^, choosing the nearest gene within 10kb of the SNP either upstream or downstream.

To put our results in context, it is of great interest to compare them to findings obtained from traditional iQT-based GWASs. To do this, we searched two leading databases for such findings: the GWAS Catalog and the Oxford Brain Imaging Genetics (BIG) Server (https://open.win.ox.ac.uk/ukbiobank/big40/)^20^. In GWAS Catalog, we use “brain measurement” (EFO ID: EFO_0004464) to query. A total of 1,742 SNPs were found to be associated with various brain iQTs (p-value threshold 1 × 10^-5^). For the BIG server, we included all the 4,323 associated (using the default p-value threshold 1 × 10^-7.5^) SNPs.

### 9. Saliency map construction

We visualized CNN model attention using the *tf_keras_vis* package (v0.8.1)^55^. This package provides various options for visualization, and we chose the Grad-CAM++^46^ method to visualize the regions of the input image that contributes the most to the output value. Grad-CAM++ uses a weighted combination of the positive partial derivatives of the last convolutional layer feature maps with respect to a specific class score as weights to generate a visual explanation for the corresponding class label. During model training stage for each model, we saved the best model weights and parameters. To construct the saliency maps, we first loaded the weights of best model, we then replaced the SoftMax activation function in the last layer with a linear activation function. The score function for the saliency map was defined as the class labels and the test dataset was used to construct the saliency map for each SNP. Since the saliency map varies sample by sample, we take the numerical mean of all saliency maps derived from all samples in one class to get the class-level saliency map. Finally, the saliency maps were plotted using the Python matplotlib library with the “jet” color map.

## Discussion

In this study, we present a novel strategy that enables GWAS to be conducted on high-dimensional structured data such as brain MR images. The idea is to cast the GWAS on full-frame MR images under a classification framework in which the genotypes of an SNP define the classes (referred as labels in the machine learning literature), the full-frame MR images are treated as input. Under the new framework, determining the association between a SNP and brain-related traits relies on whether a reasonably trained classifier is capable of differentiating MR images belonging to separate genotype-defined classes. Compared to traditional GWAS which is hypothesis testing-based, our new classification-based strategy has two fundamental advantages: first, it can handle ultra-high dimensional phenotypes such as images and second, it is capable of picking up complex, nonlinear relation between genotypes and phenotypes.

Given the same input data, the two strategies are designed to answer slightly different questions. Under the hypothesis testing framework, input data are assumed to be generated by two underlying distributions (often parametric), the goal is to examine the null hypothesis which states that there is no difference between the two distributions. In contrast, under the classification framework, the two groups are assumed to be distinct, the aim is to construct an effective classifier to distinguish two groups of data. The former, upon which the traditional GWAS is based, is a natural choice for univariate phenotypes, whereas the latter, handles multivariate traits much more elegantly. Using simulation, we showed that when applied to the same univariate trait, the two strategies return consistent results. Therefore, we believe the novel classification-based strategy can extend the powerful GWAS approach to multivariate phenotypes, especially, structured, ultra-high-dimensional phenotypes such as images.

A key advantage of taking the entire image of the brain as opposed to summary statistics as the phenotype is to allow the algorithm to discover important new brain traits in an unsupervised fashion, and capture global, higher-order nonlinear associations between variants and brain phenotypes. This could uncover genetic variant-brain associations that are difficult to be detected using traditional GWAS conducted on curated iQTs. To demonstrate its utility, we performed a large classification-based GWAS on brain MR images using publicly available datasets from the ADNI study^22^ including 1,009 individuals. Among the top 20 most significant SNPs, 9 of them were not known to be related to any brain ROI. Through gene-set analysis, tissue and pathway enrichment analysis, we have demonstrated that many of the top SNPs have close relationships with brain development and the neuronal system. The saliency map offer clues about the relevant locations of the brain in which SNP genotypes are associated to. Our results showed that our strategy has the potential to discover novel brain-related SNPs that have not been detected in previous studies.

Traditional GWAS is designed for univariate phenotypes, in which the difference in these phenotypes is easy to recognize and measure, such as disease/normal or high/low. However, in high-dimensional data, difference exist, but it is extremely difficult to explicitly describe or quantify. For example, using real life photos, our eyes can easily distinguish different objects in the photos, such as cats or dogs, but it is almost impossible to quantitively separate them using intensity values of the images. This is a fundamental property of high-dimensional data. Therefore, we believe it is necessary to expand conventional GWAS so that it can takes on high-dimensional data such as images. Here the key idea is to cast the association problem under a classification framework. If exposure to genetic mutation led to MR images with difference that is recognizable by a trained classifier, then we believe it is likely that there is strong association between this variant and brain-related phenotypes. Although we do not know immediately what exact phenotype is being impacted by the variant, we are nevertheless confident that the impacts of these variants on the MR images are substantial enough to be noticeable by artificial intelligence (AI). Therefore, we believe our strategy extend the definition of phenotypes, and has the potential to uncover novel genotype-phenotype associations and is complimentary to existing GWASs. Results obtained when applying our strategy on ADNI MR images support our hypothesis.

Compared to hypothesis testing-based strategy, our proposed classification-based strategy is more general, because difference in a model parameter (usually univariate) between two populations is a special case for two different populations.

Superficially, our strategy resembles phenome-wide association study (pheWAS), a study design that explore association between SNPs and a wide range of phenotypes^10^. Unlike GWASs, which screen all variants in the genome to identify associations with a phenotype, a PheWAS study focuses on a single variant, and explore its association with a large collection of phenotypes, often clinical diagnoses collected from electronic health record (EHR)^11–15^. The key difference between our strategy and PheWAS is that our method works with high-dimensional phenotypes, such as full-frame MR images as a whole. In contrast, a PheWAS analyzes multiple phenotypes one by one.

Admittedly, our method has room for improvement. First, the size of the dataset used in our study is limited. The data size is not large enough to develop deeper models to enhance its discriminant power. Our future work will involve collecting more data from different sources such as the UK Biobank to improve the performance of the models.

Second, for bi-allelic SNPs, there are three genotypes. But due to the limited sample size, and the fact that a large proportion of SNPs has modest MAF, hence sample size for the “aa” genotype may be small. This creates very imbalanced classes. In order to form more balanced classes, we decided to merge Aa and aa genotypes. Admittedly, this is a trade-off between simplicity, power and specificity. The strategy is also not limited to SNPs. For other types of variants such as copy number variation (CNV), our proposed strategy is readily applicable. Additionally, classes can be defined based on gene-based or region-based mutation profiles of rare variants with appropriate thresholding.

Third, in order to increase the size of the training data, we included multiple MRI scans from the same individual, which is admittedly undesirable, and will likely reduce the discovery power for the present study. We believe this is a temporary problem which will disappear when the sample size of the study cohort is sufficiently large.

Another key limitation of our strategy is the heavy computing cost, because nearly one million CNN models need to be trained. In order to reduce computational burden, in this study, we only consider 2D brain images. Hence not all information from the original 3D image data is utilized. For future work, more efficient CNN models such as vision transformers^56^ will be explored to handle full 3D MR images.

And finally, our strategy involves classifying close to a million different classification tasks. The ultimate goal is not to improve classification accuracy in individual tasks, but to identify which task is “classifiable”. Analogous to the multiple testing problem, we are facing a “multiple classification” problem. How to make that distinction is an interesting new problem that requires more research effort which we plan to pursue in future studies.

The new classification-based GWAS framework we introduced is rather general and can accommodate a wide variety of high-dimensional, structured phenotypes. There are many diverse complex phenotypes in the real world including other imaging modalities like positron emission tomography (PET), speech signals, and digital signals produced by medical devices such as ultrasounds, PETs, and electroencephalogram (EEG). Such complex, high-dimensional phenotypes have yet to be fully analyzed in their original form. We believe the novel classification-based association study framework can be applied to a full spectrum of high-dimensional phenotypes and will have a broad impact across multiple areas of genetics making novel discoveries between genotypes and high-dimensional phenotypes.

## Supplementary Tables

Supplementary Table 1. The full list of the simulation study results. The results were obtained by running the simulation study under different values of the mean, sample sizes.

Supplementary Table 2. Summary of the basic information of all the subjects that were included in the analysis.

Supplementary Table 3. Training time and performance of different models in the sex classification task. Three different CNN architectures were used: LeNet, ResNet50, and Xception. The performance was measured by the macro F1 and MCC of running these models on the test set.

Supplementary Table 4. The full list of the classification results of all SNPs in different three orientations.

Supplementary Table 5. The full list of the permutation test results for top 500 SNPs in the axial direction and the genomic context annotation of these SNPs.

Supplementary Table 6. The full list of the permutation test results for top 500 SNPs in the coronal direction and the genomic context annotation of these SNPs.

Supplementary Table 7. The full list of the permutation test results for top 500 SNPs in the sagittal direction and the genomic context annotation of these SNPs.

Supplementary Table 8. Phenotype-Wide Association Study (PheWAS) annotation of the top 500 SNPs in each of the three directions: axial, coronal and sagittal.

Supplementary Table 9. The overlap of the three lists of top SNPs ranked by MCC. Supplementary Table 10. The overlap of the three lists of top SNPs ranked by macro F1 score.

## Supplementary Figures

**Supplementary Figure 1. Overview of the ADNI subjects included in this study. (A)** Histogram displaying the distribution of the number of MRI scans per subject. **(B)** Histogram showing the number of subjects per age group. **(C)** Pie chart representing the proportion of diagnosis in the ADNI cohort, including Alzheimer’s disease (AD), cognitively normal (CN), mild cognitive impairment (MCI), early mild cognitive impairment (EMCI), late mild cognitive impairment (LMCI), and subjective memory complaint (SMC). **(D)** Pie chart illustrating the proportion of male and female individuals in the ADNI cohort.

**Supplementary Figure 2. Scatter plot showing the relationship between the classification performance and the two sample t-test p-values under different simulation settings.** The x-axis is −log10 scale of the t-test p-values. The y-axis is the classification performance (MCC and macro F1). N = 3000, 4000, 5000, 6000, 7000. The dashed line is the GWAS threshold of 5e-8. **(A)** Dots are colored by the difference between the mean of the two groups. **(B)** Dots are colored by dfference in the proportion of the majority class.

**Supplementary Figure 3. Genomic annotation of the top 500 SNPs from classification-based GWAS conducted on three different planes--axial, coronal, and sagittal.** Overall, 1301 unique SNPs were annotated. **(A)** Venn diagram displaying overlap among the three lists of top 500 SNPs ranked by macro F1 score. **(B)** Venn diagram illustrating the overlap among the three lists of genes close to the top 500 SNPs. **(C)** Pie charts showing the proportion of variant types among all the top SNPs annotated. **(D)** Histogram of macro F1 values for all SNPs in the axial, coronal, and sagittal planes.

**Supplementary Figure 4. GTEx tissue enrichment of genes nearby (within 10kb upstream and downstream to) the top 500 SNPs in three different planes. (A)** Axia, **(B)** Coronal, **(C)** Sagittal. The top 20 tissue-specific enrichments are shown for each plane. The enrichment was calculated based on the GTEx data with online tool: https://maayanlab.cloud/Enrichr/.

