## Supplementary figures and images for "A novel classification framework for genome-wide association study of whole brain MRI images using deep learning"

### Supplementary Figure 1

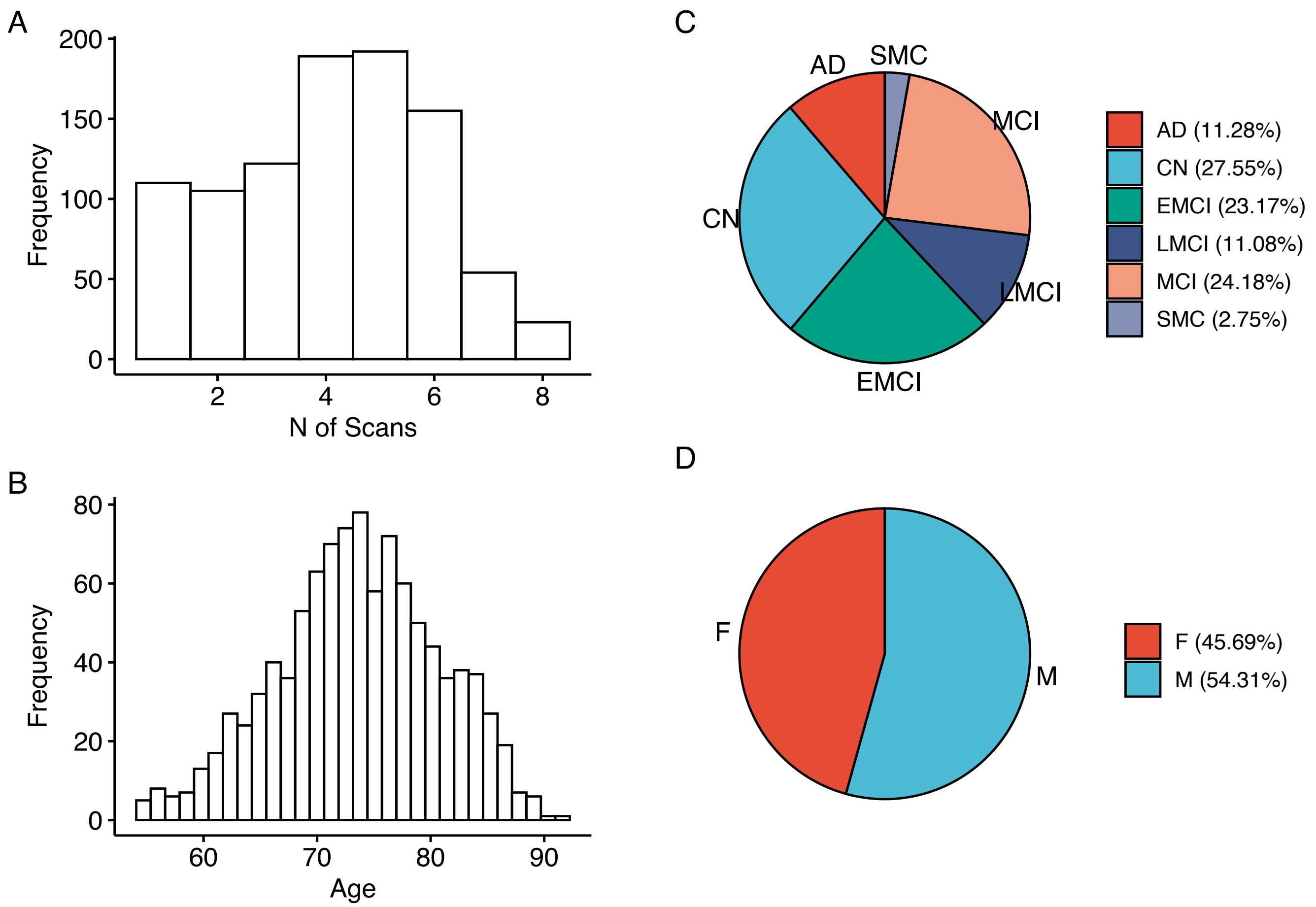

### Supplementary Figure 2

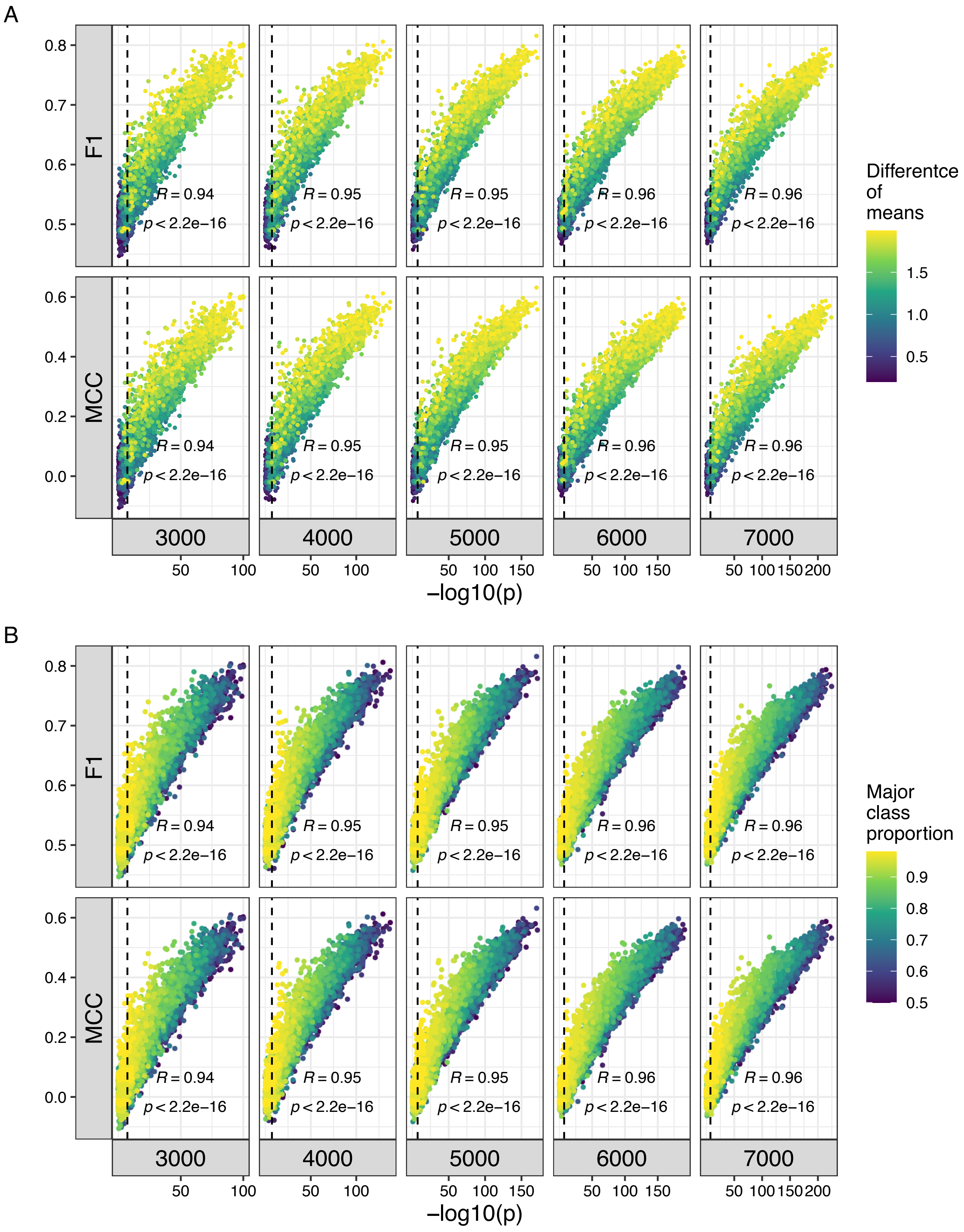

### Supplementary Figure 3

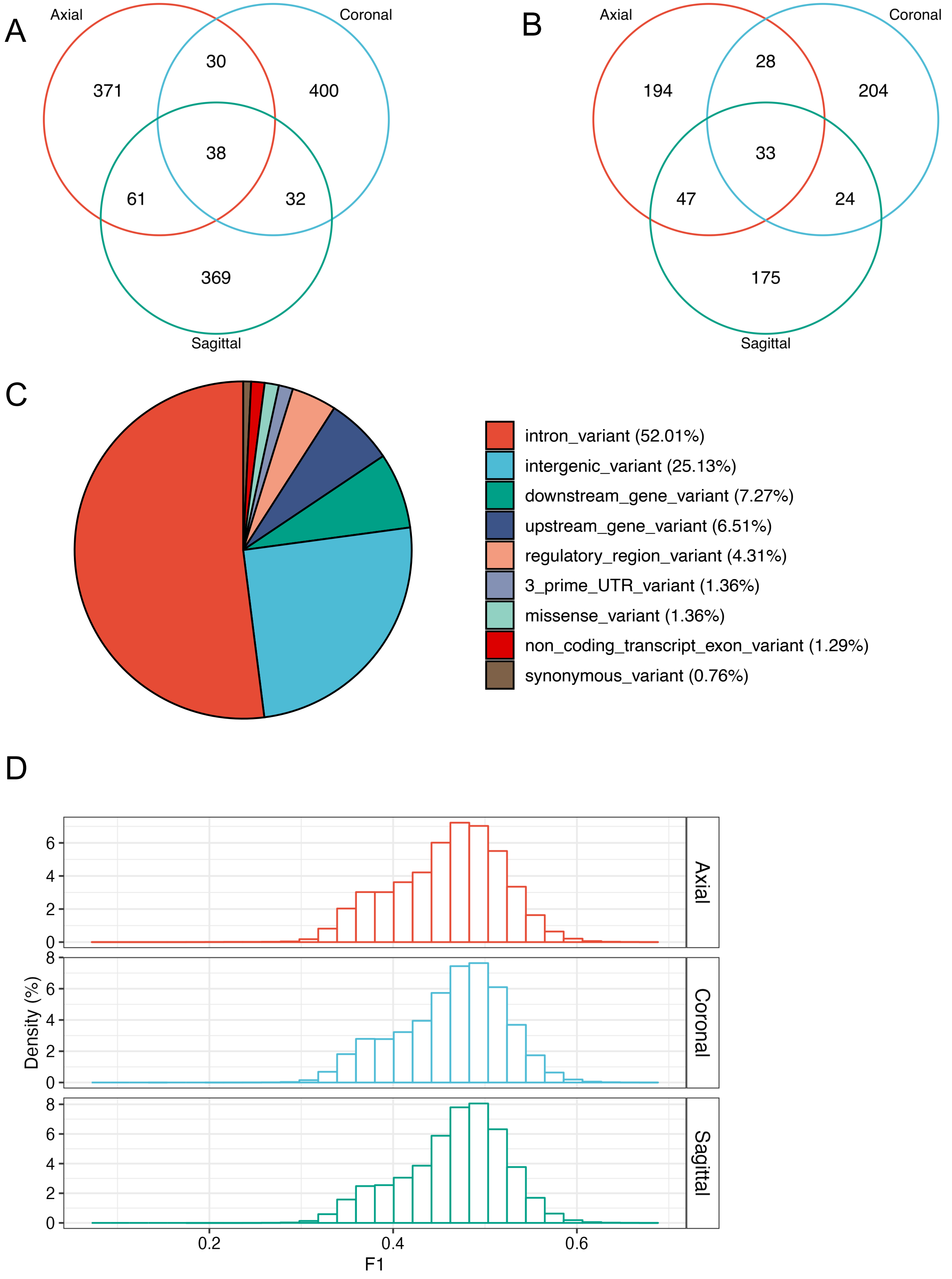

### Supplementary Figure 4

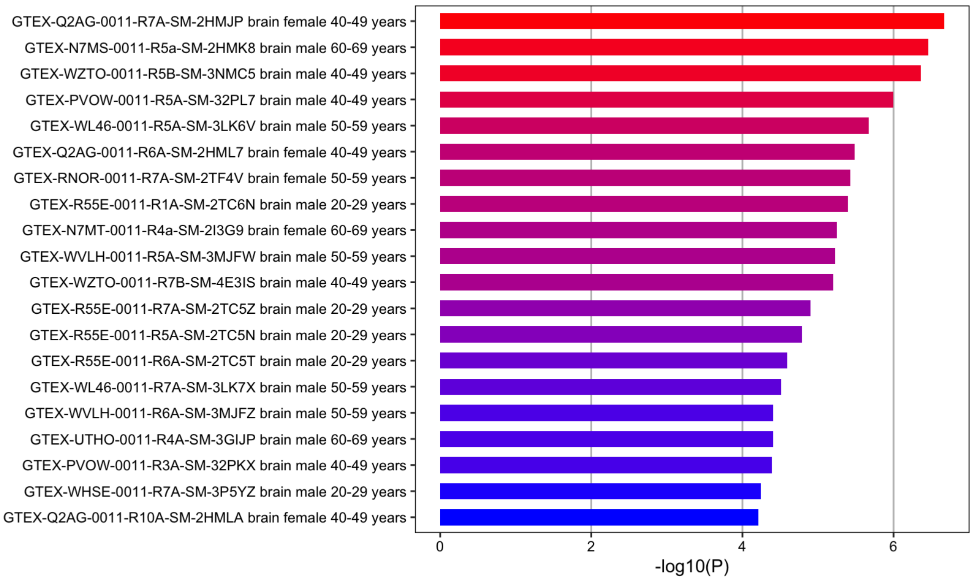

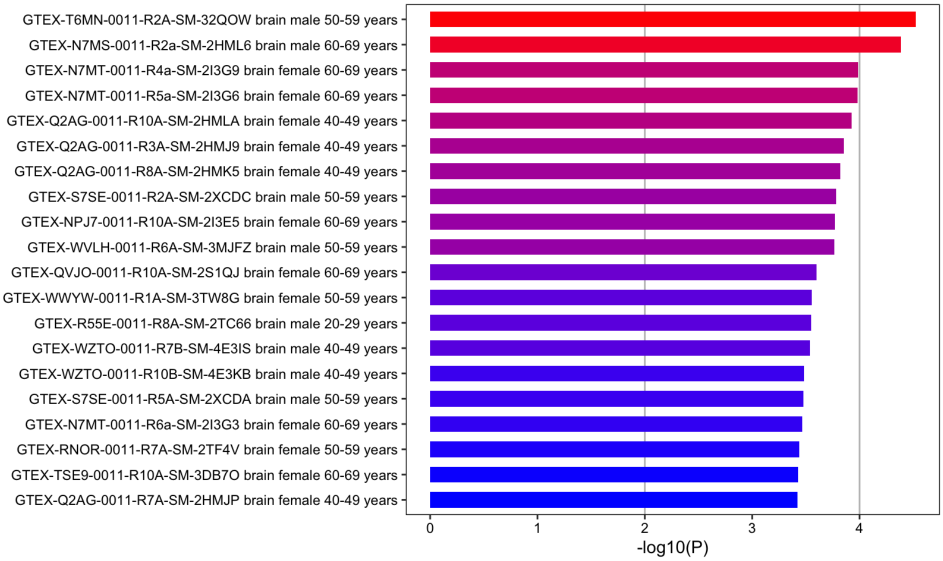

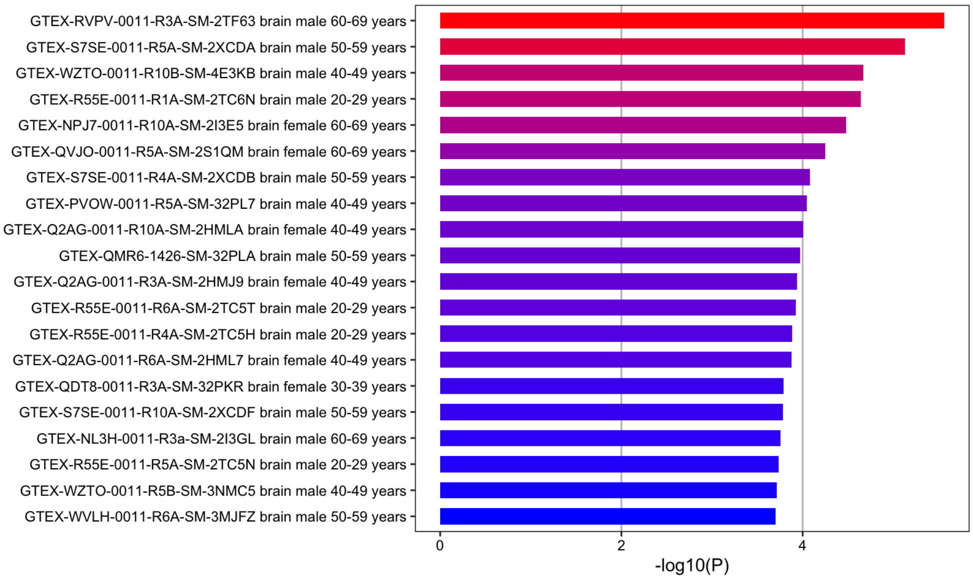
